# TEMPORAL DYNAMICS OF COMPLEMENT ACTIVATION AND OUTCOME IN PATIENTS WITH ACUTE ISCHEMIC STROKE

**DOI:** 10.1101/2025.11.21.689862

**Authors:** Yan Wang, Willeke F. Westendorp, Merijn W. Bijlsma, Michael W.T. Tanck, Wing Kit Man, Matthijs C. Brouwer, Inge A. Mulder, Jonathan M. Coutinho, Diederik van de Beek

**Author notes:** **Corresponding Author** Prof. dr. Diederik van de Beek, Department of Neurology, Academic Medical Center, Meibergdreef 9, 1105 AZ, Amsterdam, Netherlands.

## Abstract

Immune-mediated inflammatory responses exacerbate brain injury and affect the prognosis in acute ischemic stroke (AIS). We characterized longitudinal complement activation after AIS using serial plasma samples from 10 patients and 8 controls. Initial markers of classical (C1q), lectin (mannose-binding lectin [MBL]), and alternative (Factor Bb [CFBb]) pathways, as well as regulatory (Factor H [CFH]) and downstream markers (C3a, C5a and C5b-9) were measured by ELISA at 9 time points. Stroke patients showed broad complement activation with increased C1q, MBL, CFBb, C3a, C5a, and C5b-9, and reduced CFH. C1q, MBL, and CFBb peaked around day 17. MBL was higher in patients with poor outcome, CFBb was higher with good outcome, and C5b-9 in severe strokes. These findings reveal pathway-specific complement dynamics after AIS and support complement as a target for stage-specific therapy.

## INTRODUCTION

Stroke remains a leading cause of death and disability worldwide^1^. Post-stroke inflammation contributes to secondary neuronal damage and poor recovery^2^. The complement system is a key component of the innate immune system that can be activated by tissue damage and ischemia (Figure 1). The classical, lectin, and alternative complement pathways converge on C3 and C5 cleavage, generating inflammatory mediators and the membrane attack complex (C5b-9). While complement activation aids in clearing ischemic tissue, its dysregulation can exacerbate vascular and neuronal injury^3^. Experimental studies have demonstrated complement deposition in ischemic brain tissue, and complement inhibition reduces infarct size and inflammation in animal models^4,5,6,7^. Despite this evidence, the temporal evolution of complement activation after acute ischemic stroke (AIS) and their relationship with clinical characteristics and outcomes remain unclear. We therefore investigated longitudinal changes in plasma complement components and their associations with stroke severity, infarct volume, age, and functional outcome.

**Figure 1.**
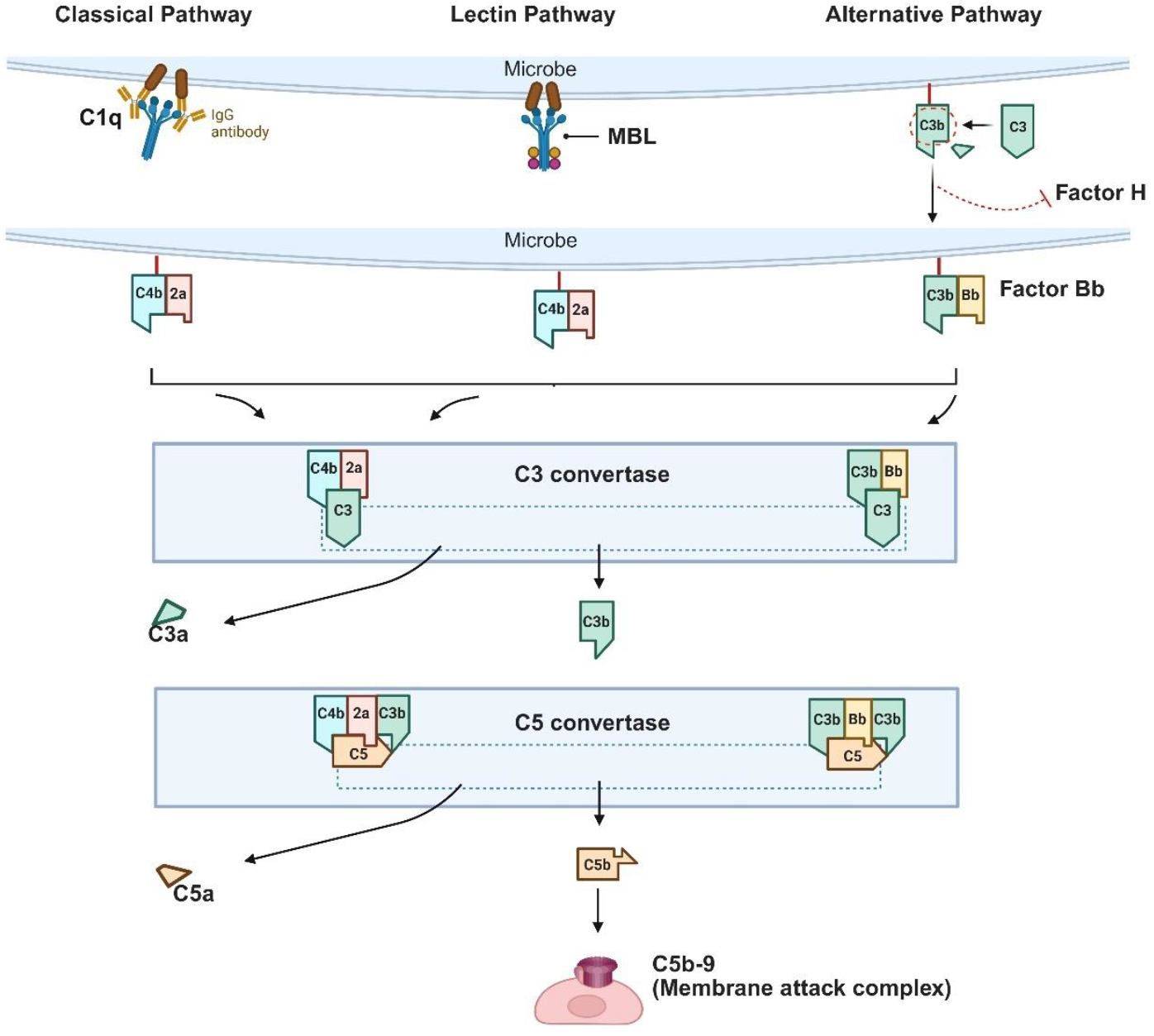
Diagram of the classical, lectin, and alternative pathways of complement activation. The initiating molecules C1q and mannose-binding lectin (MBL) trigger the classical and lectin pathways, respectively, whereas Factor B (and its active fragment, Factor Bb) participates in the alternative pathway. Factor H acts as a key regulator by competitively binding C3b, thereby dissociating or preventing the formation of the alternative pathway C3 convertase (C3bBb). C3 and C5 convertases cleave their substrates to generate the inflammatory mediators C3a and C5a. Subsequent assembly of complement components leads to the formation of the C5b-9 (membrane attack complex), which inserts into target cell membranes and cause lysis or damage.

## METHODS

We analyzed data from the TAPAS study (The Temporal Cellular Landscape of the Adaptive Immune System in Patients with Acute Stroke), a prospective case-control study conducted at Amsterdam UMC between April 15, 2021, and March 1, 2022. Ten patients with AIS and 8 age- and sex-matched controls were included. Patient inclusion criteria were ≥ 18 years, had hemispheric large-vessel ischemic stroke with symptom onset < 48 hours, the National Institutes of Health Stroke Scale (NIHSS) score ≥ 1, and were hospitalized. Exclusion criteria included immunosuppressive medication, HIV, lymphoproliferative or autoimmune disease, infection requiring antibiotics at admission, or imminent death. The study was approved by the local ethics committee, and written informed consent was obtained from all participants or their representatives.

For AIS patients, blood was collected at nine time points (days 1, 3, 7, 10, 14, 17, 21, 24, 28) and once on day 1 for control subjects. Plasma concentrations of seven complement components, initial markers of classical (C1q), lectin (mannose-binding lectin [MBL]), and alternative (Factor Bb [CFBb]) pathways, markers as well as regulatory (Factor H [CFH]) and downstream markers (C3a, C5a and C5b-9) were quantified using commercial ELISA kits: C1q, CFBb, CFH, C3a, and C5b-9 with Quidel MicroVue ELISA kits (#A001, #A027, #A040, #A032, and #A029, respectively), MBL with the Human MBL Immunoassay (R&D Systems, #DMBL00), and C5a with the BD OptEIA Human C5a ELISA Kit II (BD Biosciences, #557965).

The primary clinical outcome was modified Rankin Scale (mRS) score at 90 days, categorized as favorable functional outcome (mRS score < 3) or unfavorable outcome (mRS score ≥ 3). Secondary outcome measures included 72-hour NIHSS score (mild ≤ 7 *vs*. moderate-to-severe > 7), infarct volume on MRI (≤ 50 mL *vs*. > 50 mL), and age (≤ 68 *vs*. > 68 years).

Complement biomarker levels were normalized using a rank-based empirical normal quantile transformation (ENQT) to reduce the influence of skewed distributions and extreme values. Complement trajectories were modeled using linear mixed-effects regression with patient as a random effect and restricted cubic splines for nonlinear time effects. Associations with clinical variables were tested by including main and time-interaction terms. Results are presented as means with 95% confidence interval; p < 0.05 was considered significant. Analyses were performed in R (v4.3.2).

## RESULTS

Ten patients with AIS and 8 matched controls were included (Table 1). Median age was 68.0 years in the AIS group and 69.5 years in controls, 80% and 75% were men, respectively. The mean NIHSS score among stroke patients was 10 (range 2-18). Vascular risk factors were comparable between groups.

**Table 1.**
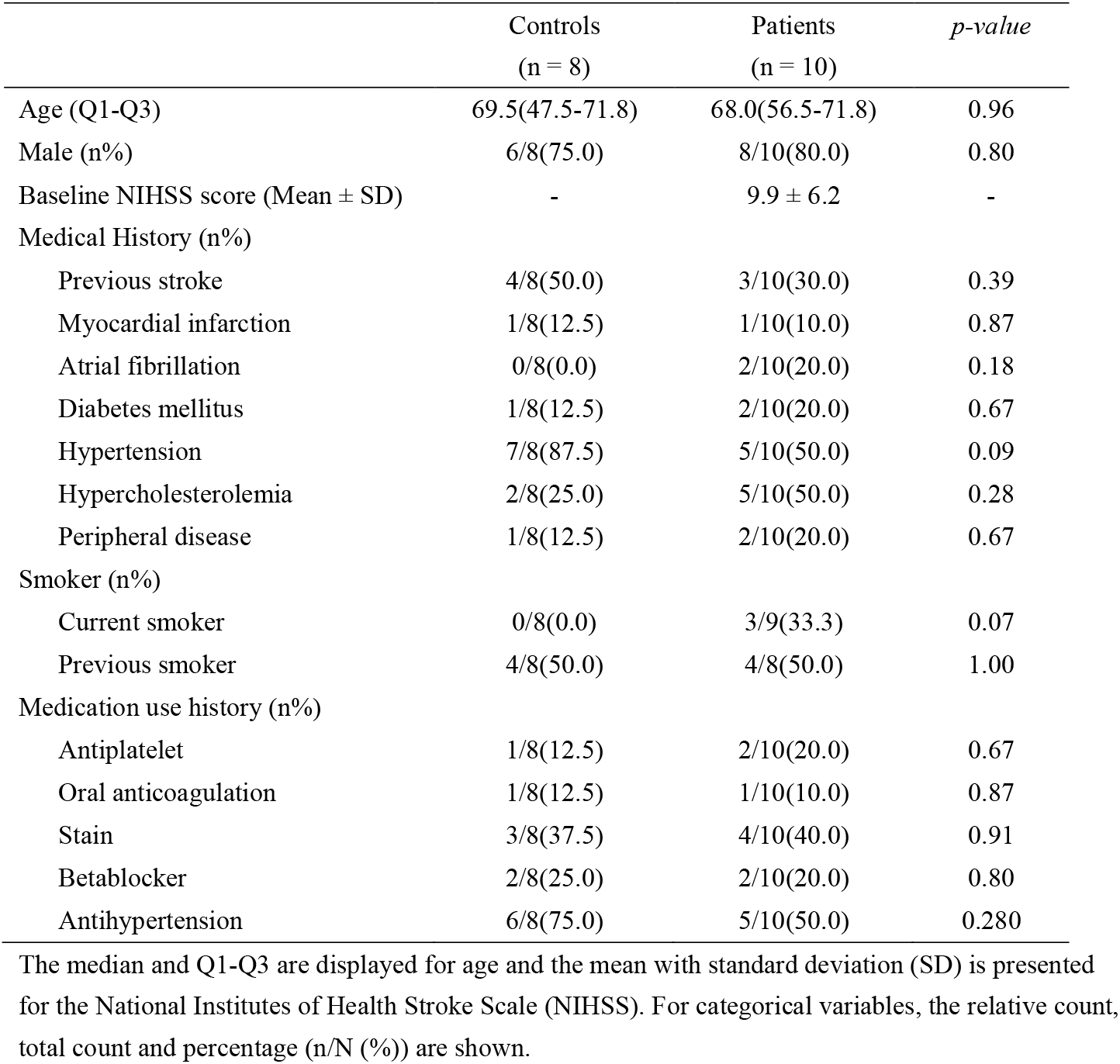
Baseline characteristics.

|  | Controls<br>(n = 8) | Patients<br>(n = 10) | *p-value* |
| --- | --- | --- | --- |
| Age (Q1-Q3) | 69.5(47.5-71.8) | 68.0(56.5-71.8) | 0.96 |
| Male (n%) | 6/8(75.0) | 8/10(80.0) | 0.80 |
| Baseline NIHSS score (Mean ± SD) | - | 9.9 ± 6.2 | - |
| Medical History (n%) |  |  |  |
| Previous stroke | 4/8(50.0) | 3/10(30.0) | 0.39 |
| Myocardial infarction | 1/8(12.5) | 1/10(10.0) | 0.87 |
| Atrial fibrillation | 0/8(0.0) | 2/10(20.0) | 0.18 |
| Diabetes mellitus | 1/8(12.5) | 2/10(20.0) | 0.67 |
| Hypertension | 7/8(87.5) | 5/10(50.0) | 0.09 |
| Hypercholesterolemia | 2/8(25.0) | 5/10(50.0) | 0.28 |
| Peripheral disease | 1/8(12.5) | 2/10(20.0) | 0.67 |
| Smoker (n%) |  |  |  |
| Current smoker | 0/8(0.0) | 3/9(33.3) | 0.07 |
| Previous smoker | 4/8(50.0) | 4/8(50.0) | 1.00 |
| Medication use history (n%) |  |  |  |
| Antiplatelet | 1/8(12.5) | 2/10(20.0) | 0.67 |
| Oral anticoagulation | 1/8(12.5) | 1/10(10.0) | 0.87 |
| Statin | 3/8(37.5) | 4/10(40.0) | 0.91 |
| Betablocker | 2/8(25.0) | 2/10(20.0) | 0.80 |
| Antihypertension | 6/8(75.0) | 5/10(50.0) | 0.280 |

The median and Q1-Q3 are displayed for age and the mean with standard deviation (SD) is presented for the National Institutes of Health Stroke Scale (NIHSS). For categorical variables, the relative count, total count and percentage (n/N (%)) are shown.

Stroke patients showed broad complement activation across the classical, lectin, and alternative pathways (Figure 2A). Concentrations of C1q, MBL, and CFBb were higher in patients than in controls, while CFH was lower. Downstream effector molecules C3a, C5a, and C5b-9 were also elevated, indicating activation of the terminal complement pathway.

**Figure 2.**
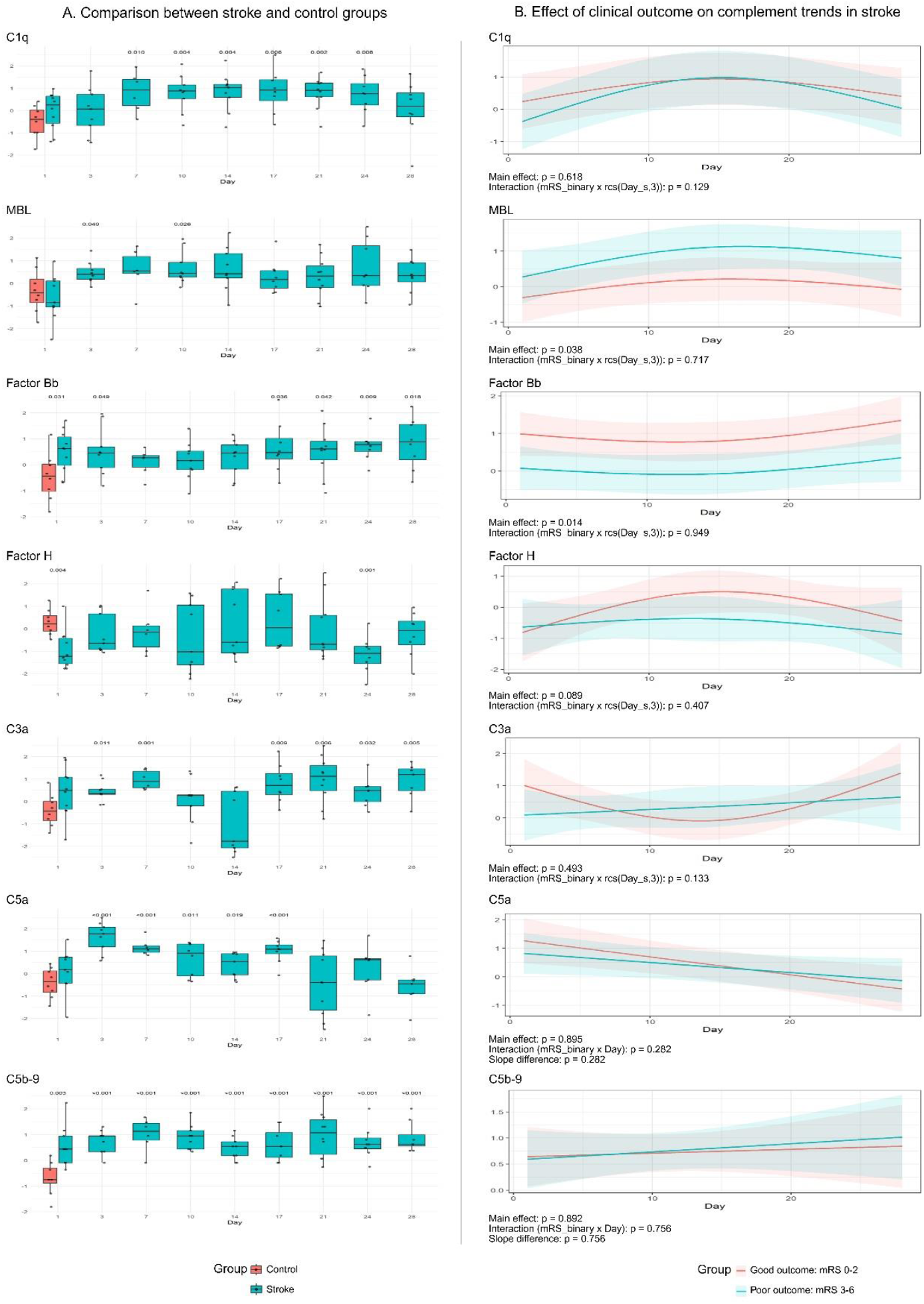
Complement component levels of acute ischemic stroke (AIS) patients and control groups (A) and outcome-related temporal trends in AIS (B). Complement levels were transformed using a rank-based empirical normal quantile transformation. (A) Red box plots: control group, green box plots: AIS group. The p-values at each time point represent comparisons between complement levels in AIS patients at the indicated time point and those in the control group on Day 1. (B) Clinical outcomes were classified according to the modified Rankin Scale (mRS) at 90 days: favorable outcome (mRS 0-2, n = 5) and unfavorable outcome (mRS 3-6, n = 5). The main effect indicates the overall difference in complement expression levels between the favorable and unfavorable outcome groups, whereas the interaction term reflects the difference in temporal trends between the two outcome groups. In the linear model, the figure also displays a comparison of the slopes between the two outcome groups. Red represents the favorable outcome group, and green represents the unfavorable outcome group.

Longitudinal modeling revealed distinct, pathway-specific trajectories (Supplementary Figure 1). C1q, MBL, and CFBb displayed significant non-linear trajectories, with C1q and MBL showing a gradual increase up to day 17, followed by a subsequent gradual decline, while CFBb exhibited an initial downward trend followed by a secondary gradual rise after day 17. CFH and C3a followed similar but non-significant nonlinear trends. In contrast, C5a declined steadily, whereas C5b-9 gradually increased throughout the 28-day follow-up. These findings suggest early engagement of the classical and lectin pathways, followed by sustained activation of the terminal complement pathway.

Complement dynamics varied with functional outcome (Figure 2B), and age, stroke severity, infarct volume (Supplementary Figures 2-4). Five patients achieved a favorable outcome (mRS 0-2) and five an unfavorable outcome (mRS 3-6) at 90 days. MBL concentrations were higher (p = 0.038) and CFBb lower (p = 0.014) in patients with unfavorable outcomes. C1q levels were higher in younger patients (≤ 68 years) compared with older patients (p = 0.025), and changed significantly over time (p = 0.003), with the decline after 17 days being slower in younger patients. Other components showed no age-related differences. A significant interaction between stroke severity and time was observed for C1q levels (p = 0.028), as severe stroke patients had lower initial C1q levels, but these rose quickly and became similar later on. Patients with mild stroke (NIHSS ≤ 7) had higher CFBb levels (p = 0.003), while those with moderate-to-severe stroke (NIHSS > 7) had higher C5b-9 concentrations (p = 0.035). C1q showed a significant time-by-volume interaction (p < 0.001), with lower early but higher late-phase levels in patients with large infarcts (> 50 mL).

## DISCUSSION

In this longitudinal study, we characterized plasma complement dynamics after AIS and their associations with clinical outcomes. Complement activation was broad, involving all three initiation pathways, yet each displayed distinct temporal patterns. The classical and lectin pathways were associated with greater stroke severity and worse recovery, whereas alternative pathway activity appeared linked to less severe stroke and favorable outcome. These findings suggest that pathway-specific complement activation may exert opposing effects during the course of ischemic injury and repair.

Our findings extend prior evidence that complement activation contributes to ischemic injury. Elevated C1q has been associated with infarct size and NIHSS severity^8^. Our longitudinal data further suggest stage-specific classical-pathway activation: potentially harmful in the early phase, but involved in repair later. MBL, which triggers the lectin pathway, was similarly associated with poor outcome, supporting sustained lectin-pathway activation as a contributor to secondary damage^9,10^. Although elevated MBL has been reported in AIS patients on admission^11^, another study suggested that acute inflammation has a limited immediate effect on MBL levels, indicating that its elevation may instead reflect an individual’s baseline status or a hepatic acute-phase response^10^. Previous reports have linked high MBL levels to larger infarcts and unfavorable recovery, consistent with our findings.

The alternative pathway CFBb showed a different pattern: its levels were highest in mild strokes and patients with favorable outcomes. This may reflect its role in amplification rather than initiation of complement activation. A proteomic analysis of 32 stroke patients and 29 controls identified CFBb among 10 upregulated proteins^12^, indicating early alternative-pathway activation. The concurrent decline of factor H, the main inhibitor of the alternative pathway, likely represents regulatory consumption in response to ongoing activation. Together, these data suggest that the alternative pathway may contribute to resolution and repair when balanced.

C3a and C5a, central nodes of the complement pathways, are closely associated with stroke prognosis. Higher baseline C3 levels predict poor 3-month outcomes and mortality in AIS^13,14^. Experimental models have shown that C3 and C5 activation increase blood-brain barrier permeability and microglial activation, leading to neuronal loss^15,16,17^. C5a activates microglia and astrocytes, inducing TNF-α and IL-6 production and amplifying inflammation^18^. In our cohort, C5a declined over time, suggesting transient activation in the acute phase. In severe COVID-19, C5a inhibition with vilobelimab reduced systemic inflammation and improved coagulation profiles and outcome in critically ill patients^19,20^. This, combined with evidence from infectious and systemic inflammatory diseases^21,22^, highlights C5a as a promising target for early intervention to limit inflammatory injury. The membrane attack complex (C5b-9) remained elevated throughout follow-up and correlated with stroke severity, indicating persistent terminal-pathway activation and endothelial injury^23,24^.

Our study is exploratory and limited by sample size. Exploratory analyses without multiple-comparison correction may increase the risk of type I error. Nonetheless, consistent temporal and outcome-related patterns across pathways underscore complement’s role in stroke pathophysiology. In summary, complement activation after AIS is dynamic and pathway-specific. Early inhibition of C5a signaling or modulation of lectin-pathway activity may represent rational, stage-specific therapeutic approaches to improve recovery after ischemic stroke.

## Non-standard Abbreviations and Acronyms

AIS: Acute Ischemic Stroke
mRS: Modified Rankin Scale
NIHSS: National Institutes of Health Stroke Scale
MBL: Mannose-Binding Lectin
CFBb: Complement Factor Bb
CFH: =Complement Factor H
TNF-α: Tumor Necrosis Factor-alpha
IL-6: Interleukin-6
ENQT: Empirical Normal Quantile Transformation

